# Standardized biogeographic grouping system for annotating populations in pharmacogenetic research

**DOI:** 10.1101/384016

**Authors:** Rachel Huddart, Alison E. Fohner, Michelle Whirl-Carrillo, Genevieve L. Wojcik, Christopher R. Gignoux, Alice B. Popejoy, Carlos D. Bustamante, Russ B. Altman, Teri E. Klein

**Affiliations:** Department of Biomedical Data Science, Stanford University, Stanford, CA 94305, USA; Department of Epidemiology, University of Washington, Seattle, WA 98195, USA; Division of Bioinformatics and Personalized Medicine, and Department of Biostatistics, University of Colorado, Anschutz Medical Campus, Aurora, CO 80045, USA; Stanford Center for Integration of Research on Genetics and Ethics, Stanford, CA 94305, USA; Department of Genetics, Stanford University, Stanford, CA 94305, USA; Department of Biomedical Engineering, Stanford University, Stanford, CA 94305, USA; Department of Medicine, Stanford University, Stanford, CA 94305, USA; These authors contributed equally to this work

## Abstract

The varying frequencies of pharmacogenetic alleles between populations have important implications for the impact of these alleles in different populations. Current population grouping methods to communicate these patterns are insufficient as they are inconsistent and fail to reflect the global distribution of genetic variability. To facilitate and standardize the reporting of variability in pharmacogenetic allele frequencies, we present seven geographically-defined groups: American, Central/South Asian, East Asian, European, Near Eastern, Oceanian, and Sub-Saharan African, and two admixed groups: African American/Afro-Caribbean and Latino. These nine groups are defined by global autosomal genetic structure and based on data from large-scale sequencing initiatives. We recognize that broadly grouping global populations is an oversimplification of human diversity and does not capture complex social and cultural identity. However, these groups meet a key need in pharmacogenetics research by enabling consistent communication of the scale of variability in global allele frequencies and are now used by PharmGKB.

## Introduction

Interindividual variability in pharmacogenes has important consequences for drug efficacy and toxicity.(1, 2) Unlike the low frequencies of alleles that are considered actionable with respect to disease risk, pharmacogenetic variants with clinical relevance are common and, in fact, both presence and absence of variants provide valuable dosing information.(3, 4) The frequencies of many pharmacogenetic alleles vary greatly by global population, meaning that people with different ancestries can have considerably different likelihoods of carrying an allele that is associated with a particular drug response. For example, the CYP3A5*3 allele has been found at a frequency of 98% in an Iranian population but at 11% in a Ngoni population from Malawi. (5, 6) A single value for global allele frequency would fail to reflect this pattern. Presenting the differences in frequencies of pharmacogenetic alleles is important for communicating the scale of their expected impact on drug response and the degree of variation between populations. This information is invaluable for furthering pharmacogenetic research and implementation.

Many pharmacogenetic studies present allelic data for very specific populations, such as from a single country or ethnic group, which are difficult to incorporate into broader research or implementation. Literature curation and gene summaries, such as those from the Pharmacogenomics Knowledgebase (PharmGKB: www.pharmgkb.org), must group these specific populations when annotating pharmacogenetic studies to allow users to easily compare information from multiple studies. As such, tagging studies with population group identifiers is an important component of knowledge extraction from curated literature. These population group labels then are used in aggregating and evaluating overall evidence for gene-drug associations, which eventually inform clinical implementation guidelines, such as those of the Clinical Pharmacogenetics Implementation Consortium (CPIC: www.cpicpgx.org).

Similar to other areas of biomedical research, (7) current methods for grouping global populations in pharmacogenetics are based on subjective, vague, and inconsistent geographical boundaries, or on populations that are geographically straightforward to cluster and reflect little admixture.(8–12) As an example of the issues with current grouping methods, some studies cluster participants of Egyptian descent with African populations, while others cluster them with Middle Eastern populations.(13, 14) While this discrepancy illustrates inconsistencies of geographic borders, the clustering of African-descent populations of the Americas with populations from Africa, as seen in the 1000 Genomes African (AFR) superpopulation, provides another example of challenges posed by employing a small number of categories to describe a broad spectrum of genomically diverse groups. The genetic patterns seen in American populations with African ancestry differs dramatically from populations in Africa due to admixture primarily with European and American Indian populations. (15–17) While sharing common ancestry, the recent admixture typically observed in the Americas can complicate average allele frequency estimation or, at a minimum, make these combined groupings less homogeneous.(16) These insufficient grouping systems, often ad-hoc and not fully representative evidence from population genomic studies, create a barrier to understanding and interpreting pharmacogenetic allele frequencies in a globally representative fashion.

Until July 2018, PharmGKB annotated studies using the five race categories defined by the US Office of Management and Budget (OMB): White, Black or African American, American Indian or Alaska Native, Asian, and Native Hawaiian or Pacific Islander, with an additional ethnicity OMB category of Hispanic/Latino. While PharmGKB serves as a global resource, these OMB groups are US-centric and, as socio-cultural measures of identity, lack the capacity to capture the scale of global human diversity. We also investigated the utility of the biogeographic categories employed by the Human Genome Diversity Panel - Centre d’Etude du Polymophisme Humain (HGDP - CEPH), which groups its 52 populations into Africa, Europe, Middle East, South and Central Asia, East Asia, Oceania and the Americas.(8, 18, 19) These population labels work well for the populations included in the HGDP data set, which are not located in ambiguous regions between group borders and which mostly contain populations with little admixture. However, papers curated at PharmGKB can include populations located all over the world, including in the transitional zones between HGDP geographical regions and admixed populations. This leads to ambiguity in how such populations would be grouped using HGDP categories. In conclusion, existing systems are insufficient for capturing the diversity of study populations in a replicable manner that is consistent with patterns of human genetic variation.

Therefore, we sought to define a grouping system of global populations that could be used consistently to annotate pharmacogenetic studies and relevant alleles, and could capture global human population genetic patterns. Using population genetics data sources, including the 1000 Genomes Phase 3 data release and the HGDP, we propose a simple and robust grouping pattern based on nine broad biogeographic regions that represent major geographic regions of the world (**Figure 1**). It is important to note that classifying individuals and communities into a few distinct groups with defined boundaries conflicts with our understanding of human variation, history, and social/cultural identities. *As a result, we respectfully present these groups as a tool to represent broad differences in frequencies of pharmacogenetic variation rather than as a classification of human diversity*.

**Figure 1:**
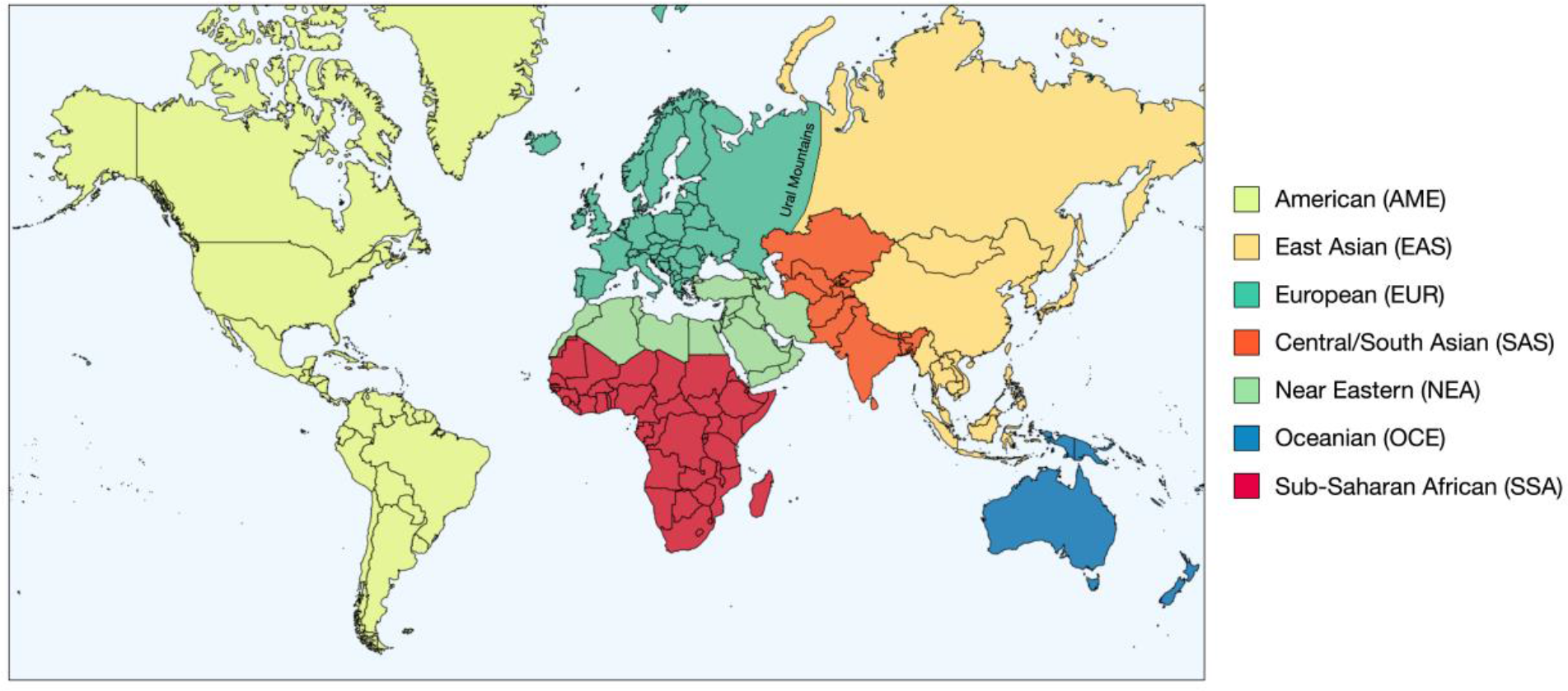
Map of geographical boundaries included in each geographical population group. Group boundaries for the seven geographical groups fall predominantly along national boundaries to aid the assignment of group membership. The two admixed groups of African American/Afro-Caribbean and Latino are not shown on this figure as the map indicates the borders of each geographical group based on the location of genetic ancestors pre-Diaspora and pre-colonization, which cannot be applied to the two admixed groups. It should also be recognized that, due to the large geographical areas covered by each group, a single group does not accurately represent the large amount of genetic diversity found in that one region.

## Results

We chose this geographic clustering pattern because geography has historically been the greatest predictor of genetic variation between human populations, with genetic distance increasing as geographic distance increases.(20) This geographic pattern aids consistency in population groupings by setting boundaries along national borders. To simplify utility, geographic boundaries between groupings are drawn predominantly along country borders, with only Russia divided into east and west along the Ural Mountains boundary due to the large size and genetic heterogeneity of the country. We intend these groups to represent peoples with a predominance of ancestors who were in the region pre-Diaspora and pre-colonization.

We have also included two admixed groups representing populations with recent gene flow between geographically-based populations and therefore, have distinct genetic patterns which are not adequately reflected by any single geographically-based group. (7) While many populations reflect a degree of admixture, we selected these two populations because they are frequently reported in pharmacogenetic studies.

We consider these nine groups sufficient to better illustrate the broad diversity in global allele frequencies, yet small enough to apply easily and to be tractable in grouping specific populations.(21–24) The groups are given below with their abbreviations.

### Geographical populations

#### American (AME)

The American genetic ancestry group includes populations from both North and South America with ancestors predating European colonization, including American Indian, Alaska Native, First Nations, Inuit, and Métis in Canada, and Indigenous peoples of Central and South America.

#### Central/South Asian (SAS)

The Central and South Asian genetic ancestry group includes populations from Pakistan, Sri Lanka, Bangladesh, India, and ranges from Afghanistan to the western border of China.

#### East Asian (EAS)

The East Asian genetic ancestry group includes populations from Japan, Korea, and China, and stretches from mainland Southeast Asia through the islands of Southeast Asia. In addition, it includes portions of central Asia and Russia east of the Ural Mountains.

#### European (EUR)

The European genetic ancestry group includes populations of primarily European descent, including European Americans. We define the European region as extending west from the Ural Mountains and south to the Turkish and Bulgarian border.

#### Near Eastern (NEA)

The Near Eastern genetic ancestry group encompasses populations from northern Africa, the Middle East, and the Caucasus. It includes Turkey and African nations north of the Saharan Desert.

#### Oceanian (OCE)

The Oceanian genetic ancestry group includes pre-colonial populations of the Pacific Islands, including Hawaii, Australia, and Papua New Guinea.

#### Sub-Saharan African (SSA)

The Sub-Saharan African genetic ancestry group includes individuals from all regions in Sub-Saharan Africa, including Madagascar.(25)

### Admixed populations

#### African American/Afro-Caribbean (AAC)

Individuals in the African American/Afro-Caribbean genetic ancestry group reflect the extensive admixture between African, European, and Indigenous ancestries(26) and, as such, display a unique genetic profile compared to individuals from each of those lineages alone. Examples within this cluster include the Coriell Institute’s African Caribbean in Barbados (ACB) population and the African Americans from the Southwest US (ASW) population, (27) and individuals from Jamaica and the US Virgin Islands.

#### Latino (LAT)

The Latino genetic ancestry group is not defined by an exclusive geographic region, but includes individuals of Mestizo descent, individuals from Latin America, and self-identified Latino individuals in the United States. Like the African American/Afro-Caribbean group, the admixture in this population creates a unique genetic pattern compared to any of the discrete geographic regions, with individuals reflecting mixed Native and Indigenous American, European, and African ancestry.

The Central/South Asian, East Asian and European groups presented here are equivalent to the 1000 Genomes South Asian (SAS), East Asian (EAS) and European (EUR) super populations, respectively. As such, we have adopted the relevant 1000 Genomes super population codes as abbreviations for each of these groups to maintain consistency. While the 1000 Genomes Ad Mixed American (AMR) super population shows complete overlap with the Latino group, we have opted to use the abbreviation LAT for this group. This removes the potential for confusion between the Latino group and the other admixed group of African American/Afro-Caribbean.

**Figure 1** illustrates the countries included in each of the seven geographical groups and removes any ambiguity of the group boundaries. As this map shows the boundaries of each group pre-colonization and pre-Diaspora, the two admixed groups, African American/Afro-Caribbean and Latino are not shown. We intend this map to be used as a guide for grouping genetic ancestral populations. Study subjects of an ancestry that is not within the geographic cluster in which they currently live will be included in the geographic cluster reflecting their ancestry. For example, South Africans of Dutch descent would be included in the European cluster rather than the Sub-Saharan African cluster. However, when lacking a clear description otherwise, the population will be included in the group that includes its home country.

This approach highlights the importance of understanding and recording detailed self-identified and self-reported race and ethnicity in the context of genetic studies. While self-reported race and ethnicity can be influenced by an individual’s social and cultural background and thus may not perfectly correlate with genetic ancestry (28), it is more reliable than assignment of race or ethnicity by another person (e.g. a healthcare professional) (29). However, it should be noted that self-reported measures can be complicated by collection processes, (30) including an incomplete selection of possible identity categories, or allowing only one selection and thus failing to capture whether an individual may identify with multiple categories or none at all (29). These classification limitations can be particularly prevalent among populations with a high degree of admixture.

To validate the genetic variability distinguished by these population groups, we conducted Principal Components Analysis (PCA) using autosomal genotype data of unrelated individuals from 1000 Genomes and HGDP. As seen in **Figure 2A**, the first two principal components (PCs) separate populations by geographic region, especially along continental boundaries, and illustrate the increasing genetic distance between populations of increasing geographic distance. As can be seen in the overlapping PC distribution of individuals of different population groups, human genetic diversity is a spectrum,(19) and therefore the geographic boundaries of these groups should be understood as an obligatory divide to create relevant groupings, with the acknowledgement that these borders are constrained by modern country borders and therefore are inherently arbitrary in geographic space.(19) However, as shown in **Figure 2B**, only a few PCs are needed to accurately predict these population clusters. Even with only 4 PCs, the minimum area under the curve (AUC) for correct cluster prediction is 97.9% for most populations using multiple logistic regression. The only outlier is the African American/Afro-Caribbean cluster, consistent with ancestral similarity to the African cluster.(15, 31) Here still, with a larger number of PCs, the AUC is above 93%, even with the observed ancestry outliers present in the 1000 Genomes African Americans in the Southwest US (ASW) population.(32) While no categorization will result in perfect prediction, given the spectrum of human diversity, the statistical validation of this clustering from broad autosomal data makes these clusters both relevant and useful for PharmGKB.

**Figure 2:**
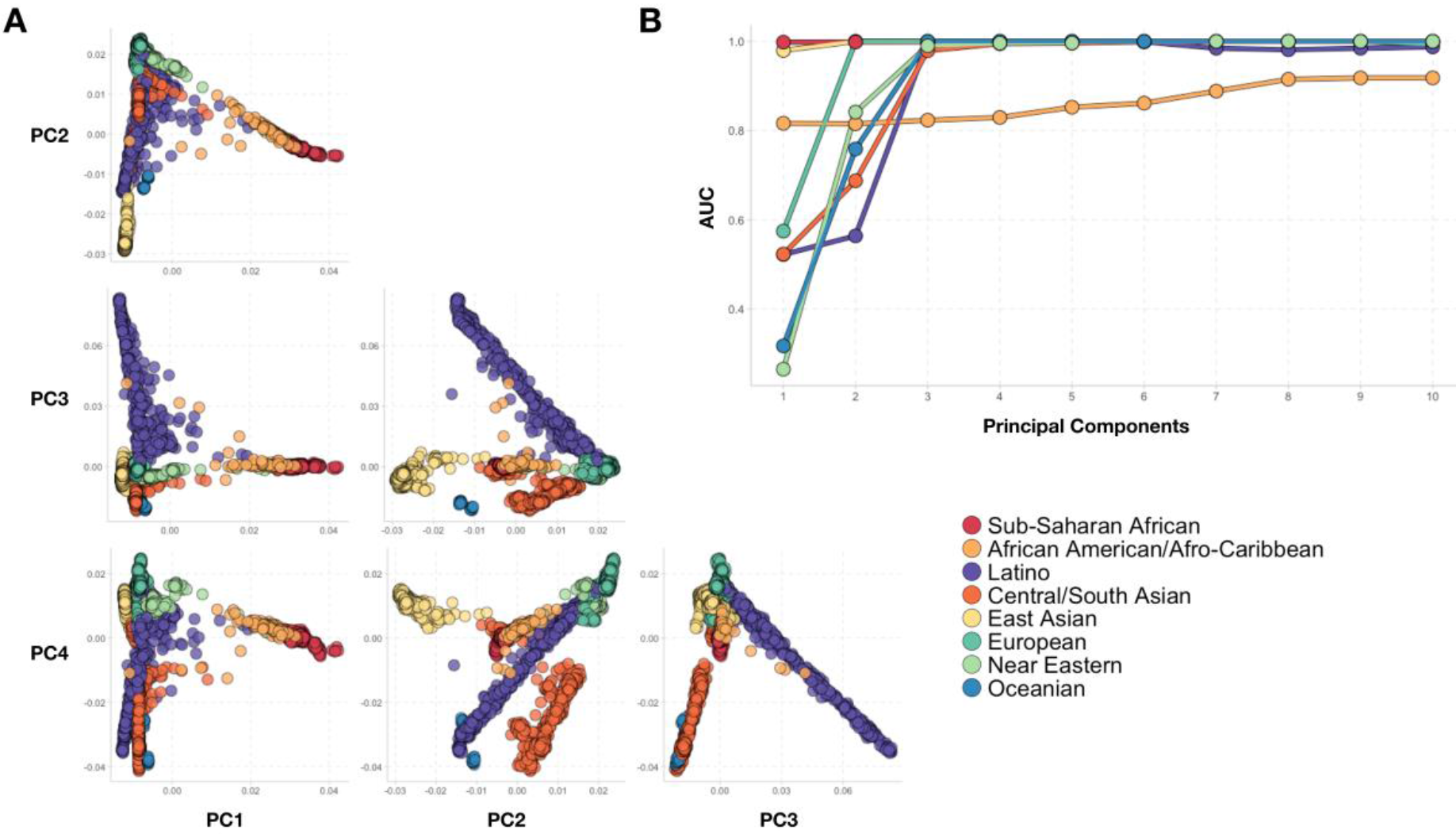
Principal component analysis comparing genetic distances of populations with close geographic proximity using 1000 Genomes participants. (A) The genetic gradient between populations is illustrated along PCs 1 vs 2 and PCs 3 vs 4, showing that, while completely discrete population boundaries are challenging, the groupings proposed here provide a statistically robust grouping. (B) AUCs of logistic regression to predict cluster membership, showing high degree of population structure. Note that, because none of the 1000 Genomes populations fall into the American (AME) group, no reference data were available to include this group in the analysis.

In **Figure 3**, we demonstrate that the groups we have selected are effective for representing the diversity of global allele frequencies in pharmacogenes. We present here the frequency of four single nucleotide polymorphisms (SNPs) with important pharmacogenetic implications. The ‘A’ allele of rs1065852 is the defining SNP of the *cytochrome P450 2D6 (CYP2D6) *10* haplotype and is also found in combination with other variants in multiple CYP2D6 haplotypes. Haplotypes containing this SNP are associated with decreased CYP2D6 activity, which has important implications for drugs that are CYP2D6 substrates, including codeine, selective serotonin reuptake inhibitors, ondansetron, and tricyclic antidepressants.(33–36) The *CYP2C9* alleles *\*2* (defined by rs1799853), *\*3* (defined by rs1057910), and *\*8* (defined by rs7900194) are associated with reduced enzyme function and therefore are associated with recommended changes to the dosing of warfarin and phenytoin, which are substrates of CYP2C9.(37, 38) Using data from the 1000 Genomes, we show the frequency of the four SNPs in these biogeographic groups. The range of frequencies between populations illustrates the importance of showing allele frequency by group in order to convey its impact on drug response globally.

**Figure 3:**
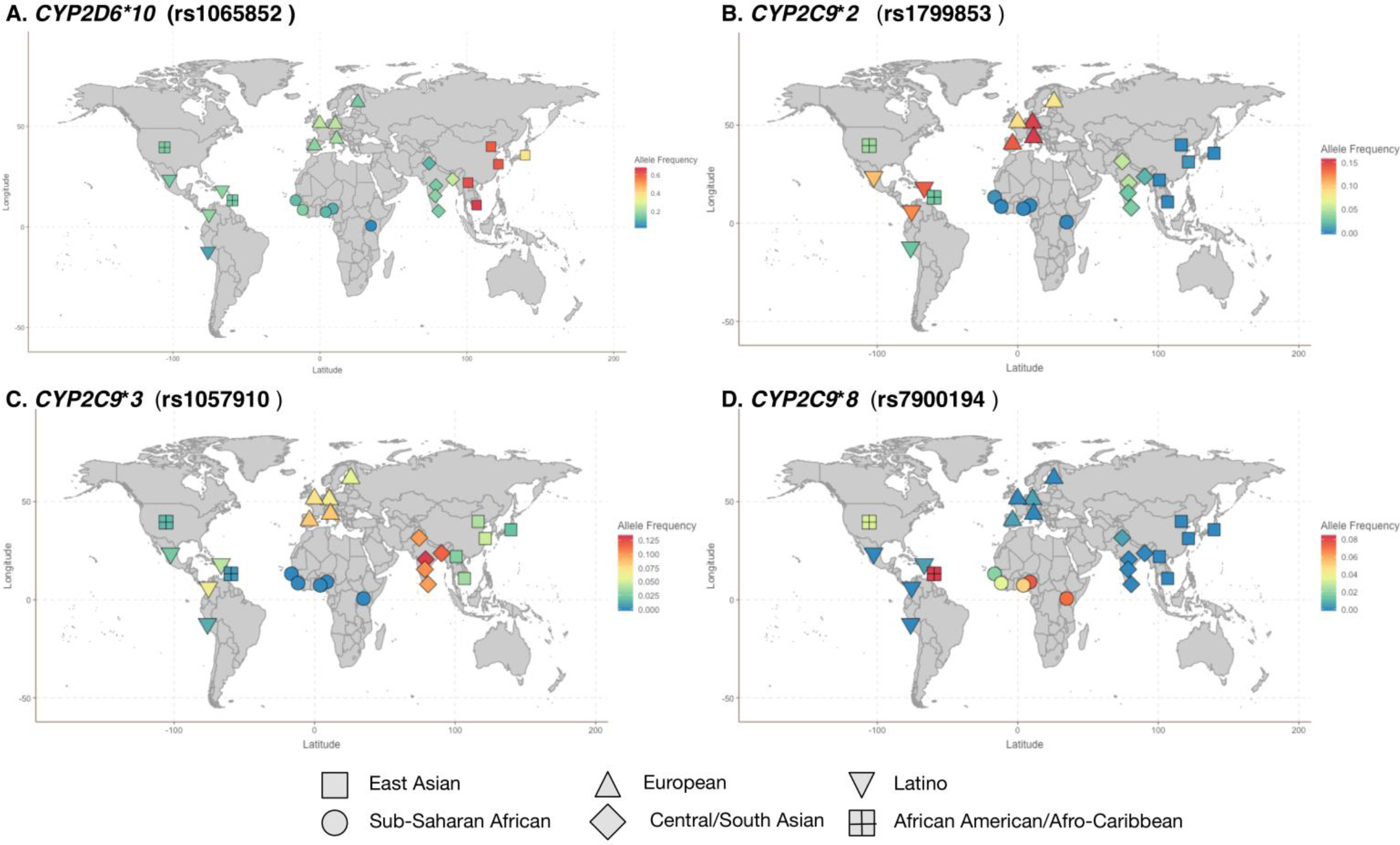
Maps illustrating how the proposed biogeographical grouping system can be used to illustrate the variability in global frequencies of key pharmacogenetic alleles. Allele frequencies from 1000 Genomes are shown across global populations for (A) CYP2D6*10, (B) CYP2C9*2, (C) CYP2C9*3 and (D) CYP2C9*8.

The SNP rs1065852 shows stark continental patterns (**Figure 3A**). The ‘A’ allele is found at high frequencies within East Asian populations, ranging from 66.2% in Vietnam (KHV) to 36.1% in Japan (JPT). This allele is less frequent in other continental populations, such as Sub-Saharan African (3.5-16.5%), European (14.6-24.7%), and Central/South Asian (10.4-25.6%).

As can be seen from the range of frequencies of the three *CYP2C9* alleles, the most common reduced function allele varies globally, with the *\*8* allele much more common in Sub-Saharan African populations (1.8-7.6%) than the *\*2* (<1%) or *\*3* (monomorphic in Africa) (**Figure 3B-D**). Conversely, the *\*8* allele is rare in European populations (<1%), while *\*2* (8.1-15.2%) and *\*3* (5.6-8.4%) are more common. Patterns such as this one can result in bias in the utility of dosing algorithms, such as the International Warfarin Pharmacogenetics Consortium (IWPC) dosing algorithm for warfarin, which adjusts dose based on the presence of the *\*2* and *\*3* alleles but does not include the *\*8* allele.(39)

## Discussion

While individual pharmacogenetic testing (either pre-emptive or at point-of-care) remains the most effective and appropriate way to implement pharmacogenetic knowledge for the care of an individual,(40, 41) we recognize the need in clinical and genetic research for a standardized method to broadly group populations based on biogeographic region. For example, identifying populations with high frequencies of certain pharmacogenetic alleles can help to direct targeted screening when resources are constrained and inform priorities for future pharmacogenetic research.(20) However, the groups we present are large and the summary information presented should be understood as an approximation dependent on existing studies in that region, which may be limited to a few locations. As such, these groups are not suitable for use in guiding specific implementation programs; rather, they should be seen as a tool for research purposes.

It should be noted that this grouping system does have limitations. Classifying individuals into these population groups can be complicated by social and cultural identities(8, 10, 42–44) and membership of an individual within one of these population groups is inherently an imperfect surrogate for predicting the likelihood that the individual carries a particular genetic variant.(41, 45) As can be seen in the analysis of rs1065852 above, the frequency of the ‘A’ allele can vary by up to 30% between populations which are all included in the East Asian group. Furthermore, while the grouping system is based on overall genome-wide average patterns, which typically follow a clinal variation pattern correlated with geographic proximity,(8, 23, 24, 46, 47) variation in individual genes or individual populations do not always follow these gradual patterns.(9–12, 41) In an attempt to mitigate some of these limitations, we encourage researchers using this grouping system to also provide specific details regarding the geographical and racial or ethnic origins of their subjects.

Because aggregate annotations of pharmacogenetic research and summary allele frequencies are based only on available studies, additional studies are needed that include a greater diversity of populations to make pharmacogenetic research and allele frequency summaries more representative.(48) For example, the Sub-Saharan African (SSA) grouping represents a large swath of human genomic diversity, which is not adequately represented in the available data from HGDP and 1000 Genomes. Increased representation of these populations in pharmacogenetics studies may lead to the discovery of clinical differences within the larger grouping. Furthermore, large, reference genetic studies with targeted allele information, like that emerging from the Population Architecture using Genomics and Epidemiology (PAGE) study (www.pagestudy.org), may provide compelling evidence to adjust these group boundaries based on frequency patterns specific to pharmacogenetic alleles. Continued evolution of this grouping system will be key to ensuring that misclassification of individuals is kept to a minimum. However, it should be understood that some misclassification is inevitable and will only be truly avoided when every patient can access comprehensive pharmacogenetic testing.

Despite these limitations, broad population groups are needed for illustrating global diversity with respect to pharmacogenetic variation and the average predicted phenotypes in populations. These nine proposed biogeographic groups provide a consistent way to present these data based on a system that is grounded in robust data on population genetic patterns, and their introduction is particularly timely given the recent commentaries by Bonham *et al.* and Cooper *et al.* (7, 49) PharmGKB is now using these population groups in curation activities, and we recommend that these groups and accompanying map be considered the standard grouping mechanism for population pharmacogenetics. Ultimately, individual pharmacogenetic testing of all patients, regardless of ancestry, is needed to deliver truly personalized medicine. However, the population groups we present are useful for the standardized presentation of pharmacogenetic studies, global allele frequency summaries in pharmacogenetic research and broad clinical screening.

## Methods

The MVN joint callset for 1000 Genomes data Phase 3 (21) was downloaded directly form the website for downstream interpretation. For principal component analysis (PCA), we filtered sites with a MAF < 0.5% and thinned sites given windows of 100 kilobases or 10 variants and r2>0.2, resulting in 156,211 sites. PCA was performed in PLINK 1.9 (50). Forward stepwise logistic regression was subsequently performed, adding 1 PC at a time, to predict population labels in a bivariate fashion. Prediction accuracy was assessed using the AUC-ROC estimator, as included in the R package ‘epicalc.’ To make assessments transparent, we included all individuals with specific population labels, although it has been demonstrated in multiple venues that there are several known ancestry outliers within 1000 Genomes populations of the Americas (17, 32). Plots were performed in R and ggplot2.

### Study Highlights

#### What is the current knowledge on the topic?

The frequency of pharmacogenetic alleles can very significantly between different populations around the world. Grouping populations can simplify reporting of pharmacogenetic alleles but current methods used to group populations are inadequate and are applied inconsistently.

#### What question did this study address?

Can we improve how populations are grouped for the reporting of pharmacogenetic alleles?

#### What does this study add to our knowledge?

We present nine new biogeographical groups based on geographical location or recent genetic admixture for use in pharmacogenetic research. These groups have been validated using autosomal genetic data from large-scale sequencing initiatives.

#### How might this change clinical pharmacology or translational science?

These groups have already been adopted for use in curation activities at PharmGKB. It is hoped that use of these groups will become standard in pharmacogenetics research.

## Author Contributions

R.H., A.E.F., M.W-C., G.L.W., C.R.G., A.B.P., C.D.B., R.B.A. and T.E.K. wrote the manuscript; A.E.F., M.W-C., C.R.G. and T.E.K. designed the research, M.W-C., G.L.W. and C.R.G. analyzed the data.

